# Identifying Genomic Islands with Deep Neural Networks

**DOI:** 10.1101/525030

**Authors:** Rida Assaf, Fangfang Xia, Rick Stevens

## Abstract

**Background:** Horizontal gene transfer is the main source of adaptability for bacteria, through which genes are obtained from different sources including bacteria, archaea, viruses, and eukaryotes. This process promotes the rapid spread of genetic information across lineages, typically in the form of clusters of genes referred to as genomic islands (GIs). Different types of GIs exist, often classified by the content of their cargo genes or their means of integration and mobility. Various computational methods have been devised to detect different types of GIs, but no single method currently is capable of detecting all GIs.

**Results:** We propose a method, which we call Shutter Island, that uses a deep learning model (Inception V3, widely used in computer vision) to detect genomic islands. The intrinsic value of deep learning methods lies in their ability to generalize. Via a technique called transfer learning, the model is pre-trained on a large generic dataset and then re-trained on images that we generate to represent genomic fragments. We demonstrate that this image-based approach generalizes better than the existing tools.

**Conclusions:** We used a deep neural network and an image-based approach to detect the most out of the correct GI predictions made by other tools, in addition to making novel GI predictions. The fact that the deep neural network was retrained on only a limited number of GI datasets and then successfully generalized indicates that this approach could be applied to other problems in the field where data is still lacking or hard to curate.

## 1 Background

Interest in genomic islands resurfaced in the 1990s, when some Escherichia coli strains were found to have exclusive virulence genes that were not found in other strains [1, 2]. These genes were thought to have been acquired by these strains horizontally and were referred to as pathogenicity islands (PAIs). Further investigations showed that other types of islands carrying other types of genes exist, giving rise to more names such as “secretion islands,” “resistance islands,” and “metabolic islands,” since the genes carried by these islands could promote not only virulence but also symbiosis or catabolic pathways [3, 4, 5]. Aside from functionality, different names are also assigned to islands on the basis of their mobility. Some GIs are mobile and can thus move themselves to new hosts, such as conjugative transposons, integrative and conjugative elements (ICEs), and prophages, whereas other GIs lose their mobility [6, 7]. Prophages are viruses that infect bacteria and then remain inside the cell and replicate with the genome [8]. They are also referred to as bacteriophages in some literature, constituting the majority of viruses, and outnumbering bacteria by a factor of ten to one [9, 10]. A genomic island (GI) then is a cluster of genes that is typically between 10 kb and 200 kb in length and has been transferred horizontally [11].

Horizontal gene transfer (HGT) may contribute to anywhere between 1.6% and 32.6% of the genomes [12]. This percentage implies that a major factor in the variability across bacterial species and clades can be attributed to GIs [13]. Thus, GIs impose an additional challenge to our ability to reconstruct the evolutionary tree of life. The identification of GIs is also important for the advancement of medicine, by helping develop new vaccines and antibiotics [14] or cancer therapies [15]. For example, knowing that PAIs can carry many pathogenicity genes and virulence genes [16, 17, 18], researchers found that potential vaccine candidates resided within PAIs [19].

We demonstrate that the problem of predicting genomic islands computationally is an excellent candidate for transfer learning on visual representations, which alleviates the problem of the extreme limitation of available ground-truth datasets and enables the use of powerful deep learning technologies. We present a method (Shutter Island) that uses deep neural networks, previously trained on computer vision tasks, for the detection of genomic islands. Using a manually verified reference dataset, Shutter Island proved to be superior to the existing tools in generalizing over the union of their predicted results. Moreover, Shutter Island makes novel predictions that show GI features.

### 1.1 Related Work

Methods proposed for the prediction of GIs fall under two categories: those that rely on sequence composition analysis and those that rely on comparative genomics. We present an overview of some of these methods next.

*Islander* works by first identifying tRNA and transfer-messenger RNA genes and their fragments as end-points to candidate regions, then disqualifying candidates through a set of filters such as sequence length and the absence of an integrase gene [3]. *IslandPick* identifies GIs by comparing the query genome with a set of related genomes selected by an evolutionary distance function [20]. It uses Blast and Mauve for the genome alignment. The outcome heavily depends on the choice of reference genomes selected. *Phaster* uses BLAST against a phage-specific sequence database (the NCBI phage database and the database developed by Srividhya et al. [21]), followed by DB-SCAN [22] to cluster the hits into prophage regions. *IslandPath-DIMOB* considers a genomic fragment to be an island if it contains at least one mobility gene, in addition to 8 or more consecutive open reading frames with dinucleotide bias [23]. *SIGI-HMM* uses the Viterbi algorithm to analyze each gene’s most probable codon usage states, comparing it against codon tables representing microbial donors or highly expressed genes, and classifying it as native or nonnative accordingly [24]. *PAI-IDA* uses the sequence composition features, namely, GC content, codon usage, and dinucleotide frequency, to detect GIs [25]. *Alien Hunter* uses k-mers of variable length to perform its analysis, assigning more weight to longer k-mers [26]. *Phispy* uses random forests to classify windows based on features that include transcription strand directionality, customized AT and GC skew, protein length, and abundance of phage words [8]. *Phage Finder* classifies 10 kb windows with more than 3 bacteriophage-related proteins as GIs [27]. *Island-Viewer* is an ensemble method that combines the results of three other tools—SIGI-HMM, IslandPath-DIMOB, and IslandPick—into one web resource [28].

## 2 Results

No reliable GI dataset exists that can validate the predictions of computational methods [26]. Although several databases exist, they usually cover only specific types of GIs [Islander, PAIDB, ICEberg], which would flag as false positives any extra predictions made by those tools. Moreover, as Nelson et al. state, “The reliability of the databases has not been verified by any convincing biological evidence” [6]. We validate the quality of the predictions made by our method first by using metrics reported in previous studies, then by introducing novel metrics and presenting some qualitative assessments of the predictions.

In Table 1, we present the total number of GI predictions made by each tool over the entire testing dataset, which consists of 34 genomes and is described in more detail later.

**Table 1.**
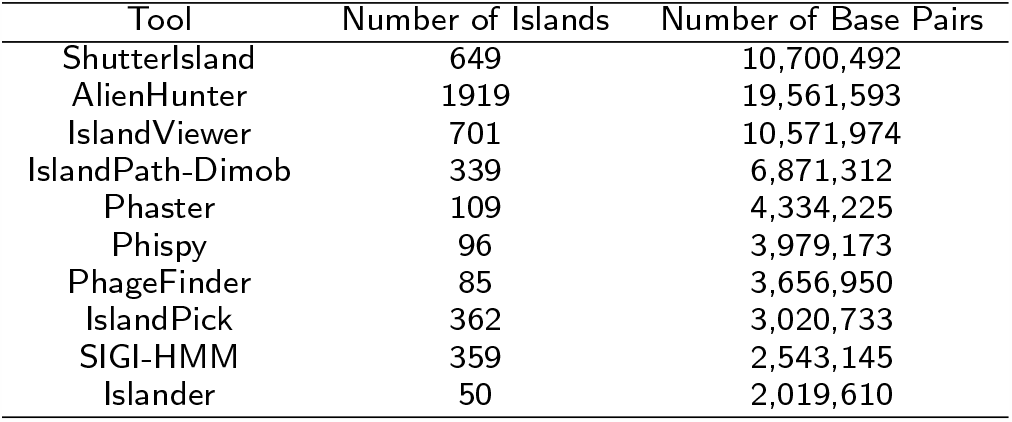
Number of islands and their total base pair value predicted by each tool over the testing genomes dataset.

We can see from Table 1 that Alien Hunter calls the most GIs, with almost double the amount called by Shutter Island and IslandViewer if measured by base pair count, and even more if measured by island count. However, it is worth mentioning that one island predicted by one tool could be predicted as several islands by another, partly due to the different length filters the tools apply. The number of predictions made by Shutter Island is close to that made by IslandViewer, which is an ensemble method combining four other tools’ predictions.

### 2.1 Validation Following Previously Accepted Methods

In this section, we present a comparison of the tools’ performance following definitions accepted by the scientific community and accepted as part of an earlier study introducing Phispy [8]. To distinguish the results presented in this section, we use *Phispy* as a pre-fix to the names of the metrics used. Namely, We refer to the metrics used as *Phispy True Positives* (PTP), *Phispy False Positives* (PFP), and *Phispy False Negatives* (PFN). Note that Phispy did not define true negatives. A true positive can be verified by the presence of phage-related genes, and a false positive by their absence. But while a region that exhibits GI features but is not predicted as a GI can be defined as a false negative, regions not showing any GI features cannot be labeled as true negatives, due to our limited understanding of GI features. Even the task of deciding the region size would not be trivial.

Tables 2 was constructed with the following definitions:

**Table 2.** True positive rate (Sensitivity) and the percentage of Phispy False Positives, as defined in the Phispy study, for predictions made by each tool over the entire testing dataset, comprised of 34 genomes.

| Tool | Sensitivity (%) | False positives (%) |
| --- | --- | --- |
| ShutterIsland | 92.4 | 29 |
| AlienHunter | 91.8 | 50.7 |
| IslandViewer | 88.2 | 30 |
| IslandPath-Dimob | 80.9 | 2.7 |
| Phaster | 73.2 | 0 |
| Phispy | 68.6 | 0 |
| PhageFinder | 65.9 | 0 |
| IslandPick | 62.5 | 44.8 |
| SIGI-HMM | 66.4 | 30.6 |
| Islander | 31.8 | 0 |

- A Phispy True Positive is a region predicted as a GI and:
  - Contains a phage-related gene, or
  - With at least 50% of its genes having unknown functions
- A Phispy False Positive is a region predicted as a GI but does not satisfy the above conditions
- A Phispy False Negative is a region with six consecutive phage-related genes that is not predicted as a GI

We followed a similar approach as the one used by Phaster to determine the presence of phage-related genes, which is looking for certain keywords present in the genes’ annotations. The set of relevant keywords can be found in the supplementary material linked to in the Additional Information section. Throughout the remainder of the paper, we refer to genes with annotations that contain such keywords as GI features.

Using the Phispy metrics defined earlier, we present the true positive rate (sensitivity) and the percentage of false positive predictions in Table 2.

Note that While some tools report 0 Phispy False Positives, they also score significantly lower on the true positive rate metric, the reason being that these tools make much fewer predictions in general.

### 2.2 Validation Using Novel Metrics

In this section, we present more general metrics to perform a more objective cross-tool comparison. Since every tool predicts a subset of all GIs, we capture the coverage of each tool across other tools’ predictions in Table 3. We omit the tools we were not able to run, and use the default parameters for all the listed tools.

**Table 3.**
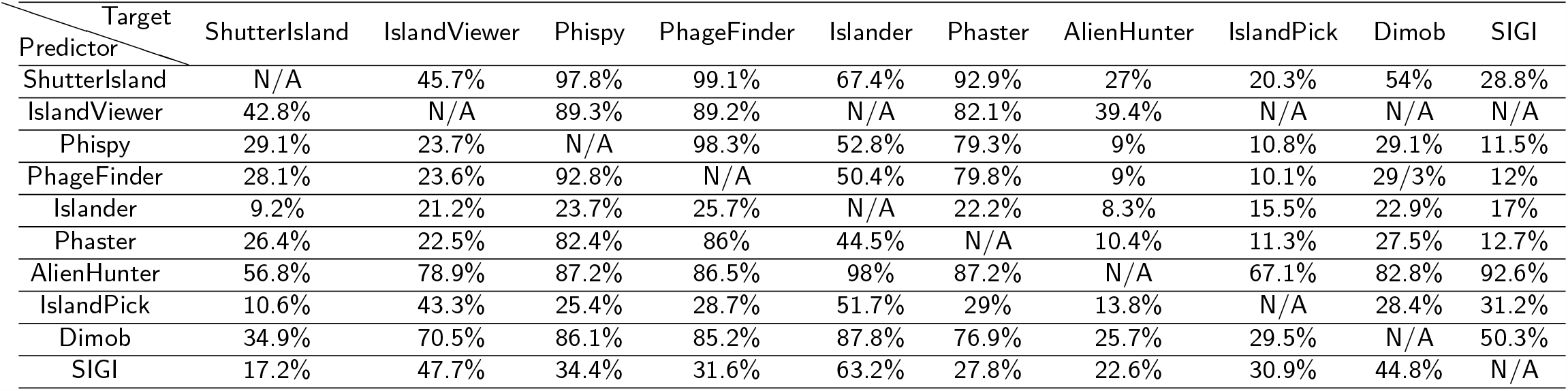
Cross-tool comparison of GI results: The percentage of GIs predicted over the testing dataset Target, that overlap with predictions made by other tools (Predictor).

| Target<br>Predictor | ShutterIsland | IslandViewer | Phispy | PhageFinder | Islander | Phaster | AlienHunter | IslandPick | Dimob | SIGI |
| --- | --- | --- | --- | --- | --- | --- | --- | --- | --- | --- |
| ShutterIsland | N/A | 45.7% | 97.8% | 99.1% | 67.4% | 92.9% | 27% | 20.3% | 54% | 28.8% |
| IslandViewer | 42.8% | N/A | 89.3% | 89.2% | N/A | 82.1% | 39.4% | N/A | N/A | N/A |
| Phispy | 29.1% | 23.7% | N/A | 98.3% | 52.8% | 79.3% | 9% | 10.8% | 29.1% | 11.5% |
| PhageFinder | 28.1% | 23.6% | 92.8% | N/A | 50.4% | 79.8% | 9% | 10.1% | 29/3% | 12% |
| Islander | 9.2% | 21.2% | 23.7% | 25.7% | N/A | 22.2% | 8.3% | 15.5% | 22.9% | 17% |
| Phaster | 26.4% | 22.5% | 82.4% | 86% | 44.5% | N/A | 10.4% | 11.3% | 27.5% | 12.7% |
| AlienHunter | 56.8% | 78.9% | 87.2% | 86.5% | 98% | 87.2% | N/A | 67.1% | 82.8% | 92.6% |
| IslandPick | 10.6% | 43.3% | 25.4% | 28.7% | 51.7% | 29% | 13.8% | N/A | 28.4% | 31.2% |
| Dimob | 34.9% | 70.5% | 86.1% | 85.2% | 87.8% | 76.9% | 25.7% | 29.5% | N/A | 50.3% |
| SIGI | 17.2% | 47.7% | 34.4% | 31.6% | 63.2% | 27.8% | 22.6% | 30.9% | 44.8% | N/A |

Table 3 shows that Alien Hunter’s predictions overlap the most with those made by other tools, which is expected given that it has the highest base-pair coverage. Shutter Island comes next and overlaps the most with three of the presented tools’ predictions. Note that while Shutter Island was trained only on the intersection of the predictions made by Phispy and IslandViewer, it generalizes and scores the highest overlap with predictions made by Phage Finder and Phaster.

Since some tools make many more predictions than do others, we used the GI features mentioned earlier to get a better idea about the quality of these overlapping predictions. In Table 4, we present the percentage of overlapping predictions that show GI features, followed by the percentage of non-overlapping predictions showing GI features. Tools that use these features to perform their classifications were omitted. We can see that on average, Shutter Island’s overlapping predictions include GI features the most. Shutter Island also misses the least predictions made by other tools that show GI features. Finally, Shutter Island has the most predictions showing GI features that are not being predicted by other tools.

**Table 4.** Quality of overlapping predictions: The percentage of GIs predicted over the testing dataset by the Target tool, that overlap with predictions made by other tools (Predictor), that include GI features — The percentage of predictions made by the Target tool but not the Predictor, that include GI features.

| Target<br>Predictor | ShutterIsland | IslandViewer | AlienHunter | IslandPick | SIGI | Average |
| --- | --- | --- | --- | --- | --- | --- |
| ShutterIsland | N/A | 91% — 64% | 87% — 47% | 89% — 31% | 87% — 36% | **89% — 45%** |
| IslandViewer | 94% — 67% | N/A | 89% — 45% | 80% — n/a | 87% — n/a | **88% — 56%** |
| AlienHunter | 74% — 70% | 66% — 60% | N/A | 73% — 21% | 71% — 42% | **71% — 48%** |
| IslandPick | 69% — 76% | 34% — 86% | 49% — 53% | N/A | 54% — 44% | **52% — 65%** |
| SIGI | 67% — 75% | 45% — 77% | 48% — 51% | 50% — 35% | N/A | **53% — 60%** |

Table 5 shows each tool’s novel predictions that do not overlap with any of other tools’, in addition to the percentage of those predictions with GI features. Alien Hunter’s unique predictions almost outnumber every other tool’s total predictions, and average 8 kbp in length. Shutter Island’s unique predictions have an average length of 14 kbp. Applying the same length cutoff threshold (8 kbp) on Alien Hunter’s unique predictions reduces them to 301 islands with a total of 3,880,000 bp, which is on par with those made by Shutter Island’s. However, a larger percentage of unique predictions made by Shutter Island exhibit GI features.

**Table 5.** The total number and base-pair count of unique predictions made by each tool over the testing genomes dataset, and the percentage of those predictions showing GI features.

| Tool | Unique GIs<br>(Count) | Unique GIs<br>(Base pairs) | Unique GIs<br>(GI features) |
| --- | --- | --- | --- |
| ShutterIsland | 280 | 3,647,377 | 65% |
| AlienHunter | 1155 | 9,583,497 | 40% |
| Phaster | 2 | 30,814 | 0% |
| Phispy | 1 | 26,890 | 100% |

Next, we show the receiver operating characteristic (ROC) curve of our classifier in Figure 1. The construction of a ROC curve requires a definition of true negative predictions. Since our classifier performs its predictions on every gene in a genome, we consider the four genes flanking each side of every query gene, and introduce the following definitions. A region is:

**Figure 1.**
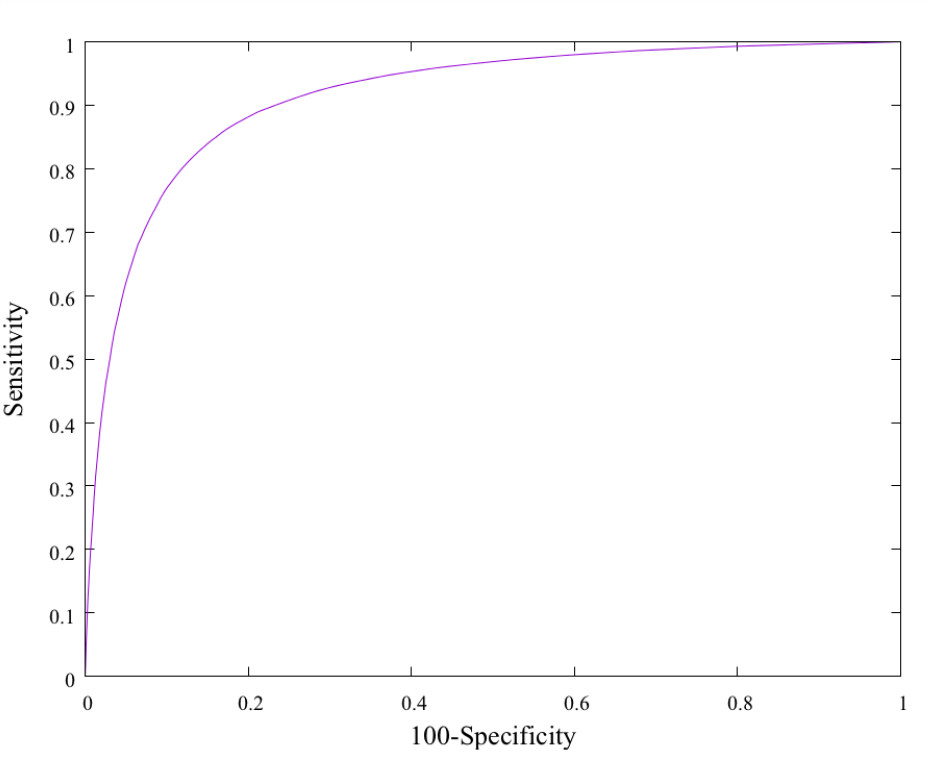
ROC curve for our genomic island binary classifier. The ROC curve plots the true positive rate as a function of the false positive rate. The greater the area under the curve is (the closer it is to the ideal top left corner point), the better.

- A true positive, if predicted as a GI and:
  - Includes a phage-related gene, or
  - Overlaps with a prediction made by another tool
- A false positive, if predicted as a GI but does not satisfy the above conditions
- A true negative, if not predicted as a GI and does not include a phage-related gene
- A false negative, if not predicted as a GI but includes a phage-related gene

### 2.3 Qualitative Assessment

To qualitatively assess the unique predictions made by Shutter Island, we present snapshots of the cargo genes typically found in these predicted regions in Figure 2, which shows that a significant number of the included genes carry GI related annotations. We also present the most common gene annotations found in the unique predictions made by Shutter Island and Alien Hunter in Figure 3. We focus on these tools since they are the ones with a significant number of unique predictions to perform the analysis on. We notice that the most frequent genes that are common to these predicted regions are either of unknown functionality or are GI-related, which adds to our confidence in these predictions.

**Figure 2.**
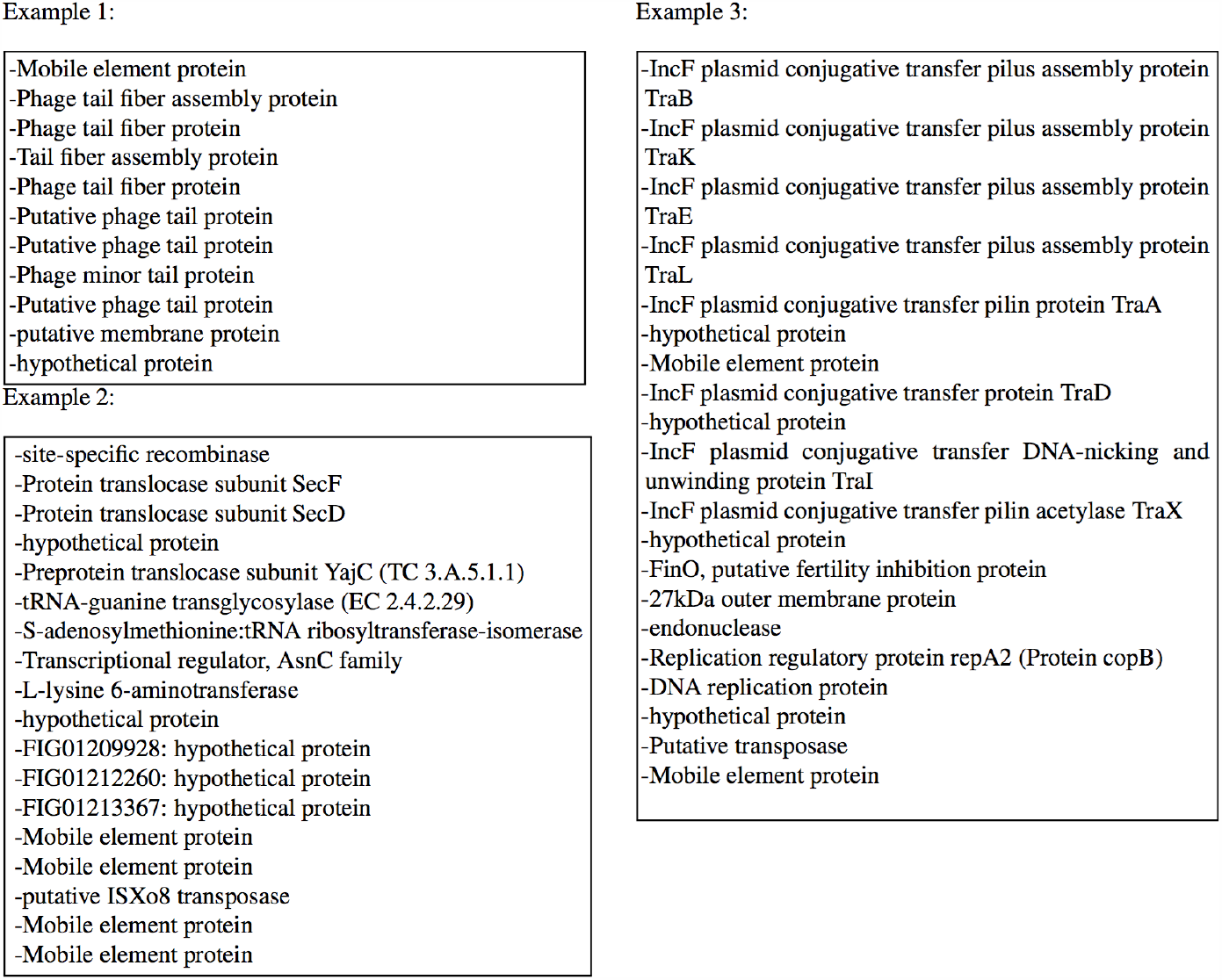
Examples of regions uniquely predicted by Shutter Island

**Figure 3.**
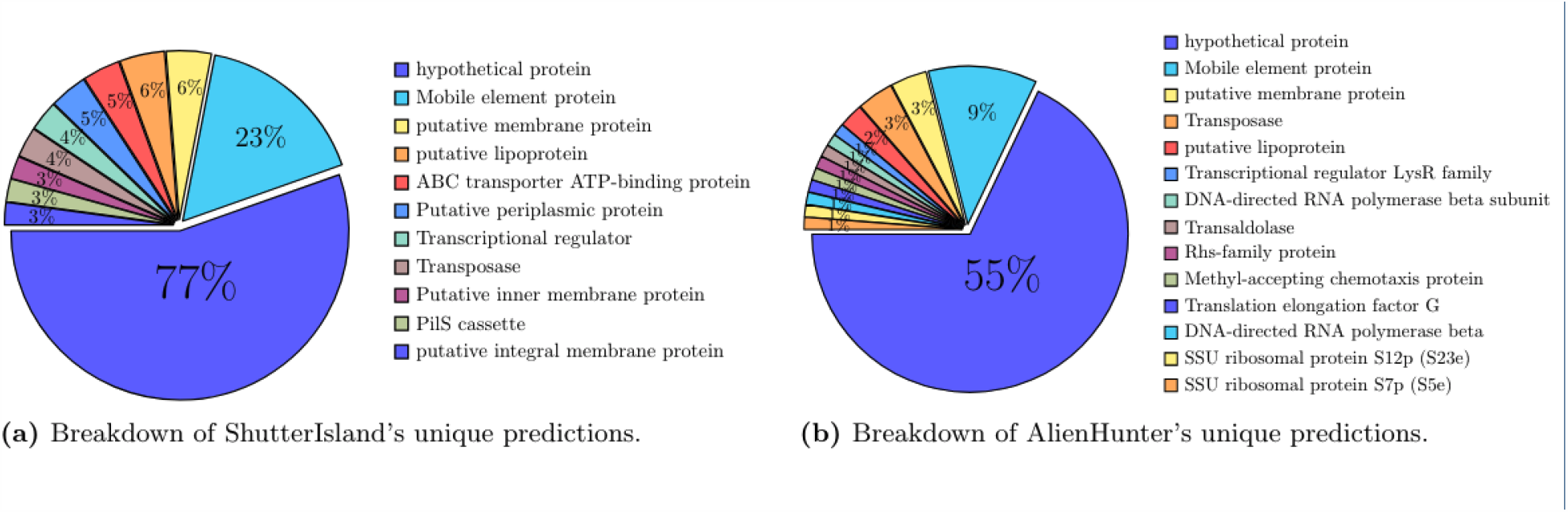
Most common gene annotations found in the unique predictions made by Shutter Island and Alien Hunter, with the percentage of unique predictions they reside in.

## 3 Methods

### 3.1 Datasets

PATRIC (the Pathosystems Resource Integration Center) is a bacterial bioinformatics resource center that we are part of (https://www.patricbrc.org) [29]. It provides researchers with the tools necessary to analyze their private data and to compare it with public data.

PATRIC recently surpassed the 200,000 publicly sequenced genomes mark, ensuring that enough genomes are available for effective comparative genomics studies. For our training data, we used the set of reference+representative genomes found on PATRIC. For each genome, our program produced an image for every non-overlapping 10 kbp window. A balanced dataset was then curated from the total set of images created. Since this is a supervised learning approach and our goal is to generalize over the tools’ predictions and beyond, we used Phispy and IslandViewer’s predictions to label the images that belong to candidate islands. IslandViewer captures the predictions of different methods, and Phispy captures different GI features. To increase our confidence in the generated labels, we labeled a genomic fragment as a GI only if it was predicted as a GI by both of these tools.

To make predictions over novel genomes, our method generates an image for every gene in the genome. Each image is then classified as either part of a GI or not. This process generates a label for every gene in the genome. A length filter of 8 kbp is then applied, so that every group of genes labeled as part of a GI that spans more than 8 kbp is reported as a single GI.

Since no reliable benchmark is available, we used the set of genomes mentioned in Phispy to test our classifier. The set consists of 41 bacterial genomes, and the authors of Phispy reported that the GIs in these genomes have been manually verified [8]. Some of the tools used in the comparison have not been updated for a while, but most of the tools had predictions made over the genomes in this testing set. We discarded the genomes for which not all the tools reported predictions over, or that were part of the training set used to train our classifier, and ended up with a total of 34 genomes, listed in the supplementary material linked to in the Additional Information section.

### 3.2 Feature Encoding Using Images

We present some of the most prominent features of genomic islands, listed by decreasing order of importance: [1, 30].

- One of the most important features of GIs is that they are sporadically distributed; that is, they are found only in certain isolates from a given strain or species.
- Since GIs are transferred horizontally across lineages and since different bacterial lineages have different sequence compositions, measures such as GC content or, more generally, oligonucleotides of various lengths (usually 2–9 nucleotides) are used [26, 31, 32]. Codon usage is a well-known metric, which is the special case of oligonucleotides of length 3.
- Since the probability of having outlying measurements decreases as the size of the region increases, tools usually use cut-off values for the minimum size of a region (or gene cluster) to be identified as a GI.
- Another type of evidence comes not from the attachment sites but from what is in between, since some genes (e.g., integrases, transposases, phage genes) are known to be associated with GIs [16]. In addition to the size of the cluster, evidence from mycobacterial phages [33] suggests that the size of the genes themselves is shorter in GIs than in the rest of the bacterial genome. Different theories suggest that this may confer mobility or packaging or replication advantages [8].
- Some GIs integrate specifically into genomic sites such as tRNA genes, introducing flanking direct repeats. Thus, the presence of such sites and repeats may be used as evidence for the presence of GIs [34, 35, 36].

Other research suggests that the directionality of the transcriptional strand and the protein length are key features in GI prediction [8]. The available tools focus on one or more of the mentioned features.

PATRIC provides a compare region viewer service, where a query genome is aligned against a set of other related genomes anchored at a specific focus gene. The service finds other genes that are of the same family as the query gene and then aligns their flanking regions accordingly, presenting the aligned pileups graphically, allowing users to visualize the genomic areas of interest. In the produced images, genomic islands appear as gaps in alignment as opposed to conserved regions. We replicated the service by implementing an offline version forked from the production user interface, which is necessary for computational efficiency and for consistency in the face of any future interface changes.

Figure 4 shows sample visualizations of different genomic fragments belonging to the two classes. Each row represents a region in a genome, with the query genome being the top row. Each arrow represents a single gene, scaled to capture the size and strand directionality. Colors represent functionality. The red arrow is reserved for the query gene, at which the alignment with the rest of the genomes is anchored. The remaining genes share the same color if they belong to the same family; they are colored black if they are not found in the query genome’s focus region. Some colors are reserved for key genes: green for mobility genes, yellow for tRNA genes, and blue for phage related genes. Figures 4(a) and 4(b) are examples of a query genome with a non-conserved neighborhood. The focus gene lacks alignments in general or is aligned with genes from other genomes with different neighborhoods from the query genome. In contrast, Figures 4(c) and 4(d) show more conserved regions, which are what we expect to see in the absence of GIs (labelled as continents in the image).

**Figure 4.**
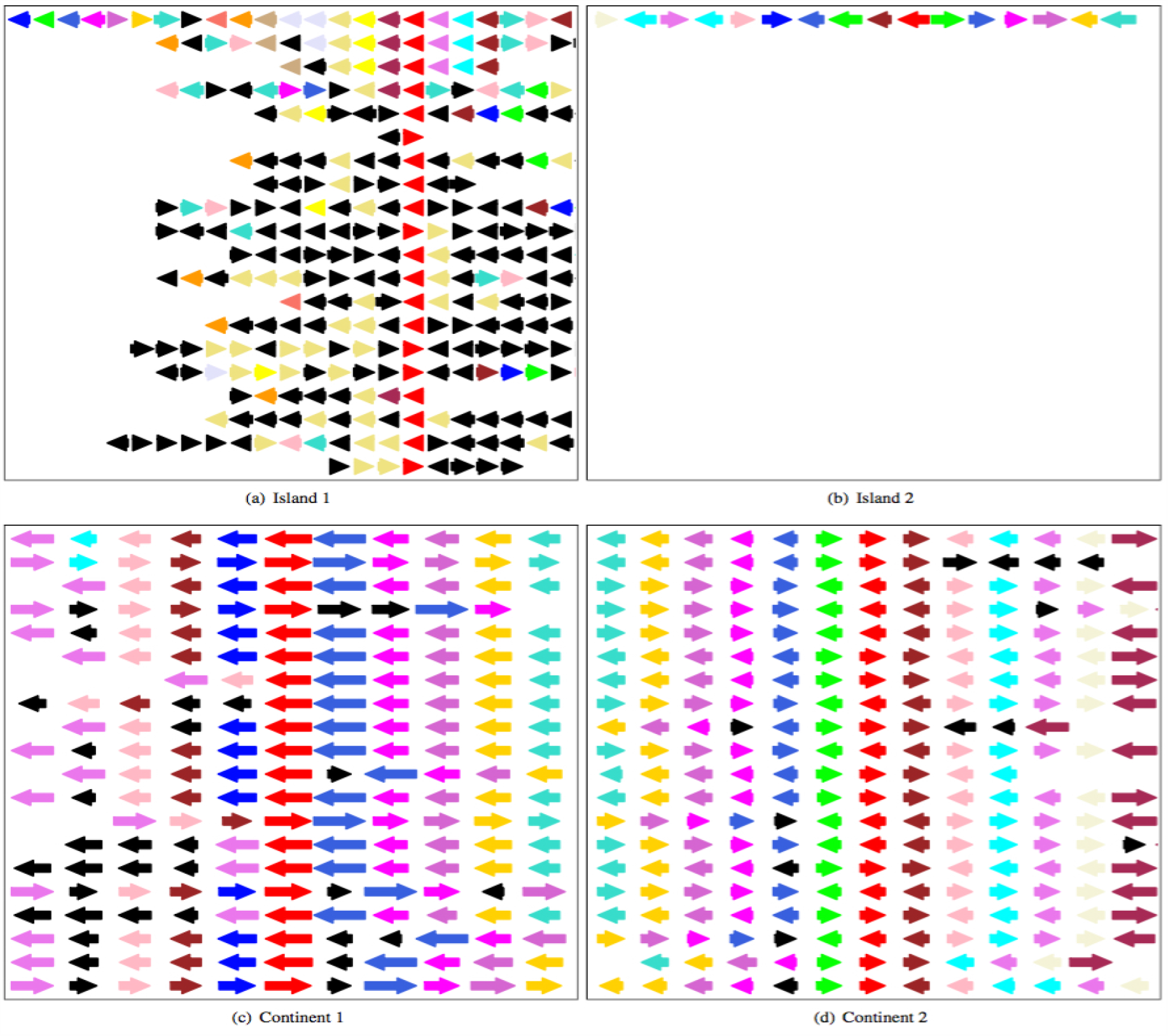
Examples of images generated using the compare region viewer. Each arrow represents a gene color coded to match its functionality. The first row is the genome neighborhood of the focus gene (red), and the subsequent rows represent anchored regions from similar genomes sorted by their phylogenetic distances to the query genome.

### 3.3 Transfer Learning

This kind of visual representation makes it easier to leverage the powerful machine learning (ML) technologies that have become the state of art in solving computer vision problems. Algorithms based on deep neural networks have proven to be superior in competitions such as the ImageNet [37]. Deep learning is the process of training neural networks with many hidden layers. The depth of these networks allows them to learn more complex patterns and higher-order relationships, at the cost of being more computationally expensive and requiring more data to work effectively. Improvements in such algorithms have been translated to improvements in a variety of domains reliant on computer vision tasks [38]. The images generated by PATRIC capture many of the most important GI features mentioned earlier—the sporadic distribution of islands, the protein length, functionality, and strand directionality—using color-coded arrows of various sizes. So, while PATRIC provides a lot of genomic data, the challenge comes down to building a meaningful training dataset. The databases available are still limited in size and specific in content, which in turn limits the ability even for advanced and deep models to learn and generalize well. Training deep models over a limited dataset puts the model at the risk of overfitting. One way around this problem is using a technique referred to as transfer learning [39]. In transfer learning, a model does not have to be trained from scratch. Instead, the idea is to retrain a model that has been previously trained on a related task. The newly retrained model should then be able to transfer its existing knowledge and apply it to the new task. This approach affords the ability to reuse models that have been trained on huge amounts of data, while adding the necessary adjustments to make them available to work with more limited datasets, adding a further advantage to our approach of representing the data visually.

In our method, we use Google’s Inception V3 neural network architecture that has been previously trained on ImageNet. Inception V3 is a 48-layer-deep convolutional neural network. Training such a deep network on a limited dataset such as the one available for GIs is unlikely to produce good results. ImageNet is a database that contains more than a million images belonging to more than a thousands categories. Thus, a model that was previously trained on ImageNet is already good at feature extraction and visual recognition. To make the model compatible with the new task, the top layer of the network is retrained on our GI dataset, while the rest of the network is left intact. Using this strategy is more favorable than starting with a deep network with random weights.

## 4 Discussion

We presented a new method, called Shutter Island, which demonstrates the effectiveness of training a convolutional neural network on visual representations of genomic fragments to identify genomic islands. In addition to using powerful technologies and extensive data, our approach may add an extra advantage over whole-genome alignment methods because performing the alignment over each gene may provide a higher local resolution and aid in resisting evolutionary effects such as recombination and others that may have happened after the integration and that usually affect GI detection efforts.

One challenge in assessing GI prediction is getting precise endpoints for predicted islands. Since different tools report a different number of islands owing to the nature of the features they use, where one island could be reported as many or vice versa, we considered a tool to predict another’s islands if any of its predictions overlap with the other tool’s predictions. We counted the percentage of base pair coverage of that other tool as represented by its predicted endpoints. This allowed us to compare overlapping islands predicted by different tools even if their coordinates did not match.

Note that when assessing the tools’ predictions, our definition of a true positive was different from the definition proposed in the Phispy study: where they define a true positive as a region with at least six phage related genes, we realize that the available datasets are much larger than what was accessible at the time of their publication, and thus the number six that was claimed to have been determined empirically may not be relevant anymore. We argue that if a region is suspected to be a GI, the mere presence of a phage related gene in that region adds to the confidence in its prediction as part of a GI. Moreover, we find that the database used by the study to determine phage functionality may be outdated, and we thus resort to certain keywords in the gene annotations generated by our system PATRIC to determine functionality. This is similar to Phaster’s validation method, whereby the presence of certain keywords (e.g, caspid) is used to verify predictions made by the tool. To identify these keywords, we scoured the literature and identified certain gene annotations that are related to GIs. Such annotations of gene identity are either directly curated by humans or reflect human assessment through exemplar-based computational propagation. We constructed a standard vocabulary of the GI-related keywords that were also in agreement with the more extensive list of keywords used by Phaster for the same purpose.

No single tool is able to detect all GIs in all bacterial genomes. Methods that narrow their search to GIs that integrate under certain conditions, such as into tRNAs, miss out on the other GIs. Similarly, not all GI regions exhibit atypical nucleotide content [40]. Evolutionary events such as gene loss and genomic rearrangement [5] present more challenges. For example, the presence of highly expressed genes or having closely related island host and donor might lead to false negatives [14]. Tools that use windows face difficulty in adjusting their size: small sizes lead to large statistical fluctuation, whereas larger sizes result in low resolution [41].

For comparative genomics methods, the outcomes depend strongly on the choice of genomes used in the alignment process. Very distant genomes may lead to false positives, and very close genomes may lead to false negatives. In general, the number of reported GIs may differ across tools, because one large GI is often reported as a few smaller ones or vice versa, making it harder to detect end points and boundaries accurately. The lack of experimentally verified ground-truth datasets spanning the different types of GIs makes point-to-point comparison across the tools extremely challenging. Moreover, different tools follow different custom-defined metrics to judge their results, typically by using a threshold representing the minimum values of features (e.g., number of phage words) present in a region to be considered a GI, which adds to the complications of validating GI predictions and comparing tools’ performances.

Our initial inspiration for representing genome features as images came from observing how human annotators work. These experts often examine the “compare region” view for a long time before they decide on the gene identity. A critical piece of information they rely on is how the focus gene compares with its homologs in related genomes. This information is cumbersome to represent in tabular data because (1) explicit all-to-all comparison is computationally expensive; (2) the comparisons need to be done at both individual gene and cluster levels including coordinates, length, and neighborhood similarities; and (3) human experts integrate all these different levels of information with an intuition for fuzzy comparison, something that is hard to replicate in tabular learning without additional parameterization or augmentation. Representing genomic features as images mitigates all three issues. Images offer a natural way to compare genes (horizontally) and clusters across genomes (vertically) with 2D convolution. The fact that the compare region view sorts genomes by evolutionary distance allows the neural network to exploit locality and place more emphasis on close genomes via incremental pooling. An additional benefit of working with images is to be able to leverage the state-of-the-art deep learning models, many of which were first developed in vision tasks and perfected over years of iterations. Google researchers have used spectrograms (instead of text) in direct speech translation [42] and DNA sequence pileup graphs (instead of alignment data) in genetic variant calling [43]. In both cases, the image-based models outperformed their respective previous state-of-theart method based on traditional domain features. Further, the low-level visual feature patterns learned in pretrained image models have been demonstrated to transfer to distant learning tasks on non-image data in several preliminary studies ranging from environmental sound classification to cancer gene expression typing [44]. Much like feature-engineering methods, casting tabular data to images encodes information in a way more amenable to learning without explicitly adding information. It can also be easily integrated with other data modalities in the latent vector representation to prevent information loss. We hypothesize that this emerging trend of representing data with images will continue until model tuning and large-scale pretraining in scientific domains start to catch up with those in computer vision.

## 5 List of abbreviations

GI: genomic island,
HGT: horizontal gene transfer,
ML: machine learning,
kbp: kilobase pair,
SNPS: single nucleotide polymorphisms

## 6 Declarations

## 7 Authors’ contributions

R.A. carried out the implementation and wrote the manuscript. R.S. and F.X. were involved in planning and supervised the work. All authors aided in interpreting the results. All authors provided critical feedback and commented on the manuscript.

## 8 Acknowledgments

We thank Dr. James J. Davis for constructive criticism of the manuscript, and Dr. Gail W. Pieper for editing the manuscript.

## 9 Funding

PATRIC has been funded in whole or in part with Federal funds from the National Institute of Allergy and Infectious Diseases, National Institutes of Health, Department of Health and Human Services [HHSN272201400027C]. Funding for open access charge: Federal funds from the National Institute of Allergy and Infectious Diseases, National Institutes of Health, Department of Health and Human Services [HHSN272201400027C]. The funding body had no direct role in the design of the study nor the collection, analysis, and interpretation of data and in writing the manuscript.

## 10 Competing interests

The authors declare that they have no competing interests.

## 11 Availability of data and materials

The datasets generated and/or analyzed during the current study are available in the GitHub repository, https://github.com/ridassaf/ShutterIsland.

## 12 Ethics approval and consent to participate

Not applicable.

## 13 Consent for publication

Not applicable.

## Notes

### Competing Interest Statement

The authors have declared no competing interest.

